# Pan-cancer RNA editing activity reveals complex editing functions and cancer immunotherapy biomarkers

**DOI:** 10.64898/2026.01.07.698309

**Authors:** Mengbiao Guo, Yuanyan Xiong

## Abstract

Pervasive RNA editing in human diversifies the transcriptome and proteome. However, biological functions of most RNA editing sites remain to be uncovered. Here, we developed a computational framework (iPEAPR) to investigate editing functions in various pathways and regulatory elements, by using only summary information of millions of editing sites, in >5,000 cancer samples and in >2,500 normal samples, and large numbers of manually curated functional annotations. We observed heterogeneous editing activities across features. Surprisingly, enhancers were among features showing the highest editing activities. Editing of epithelial-mesenchymal transition was the most significantly associated with patient survival. Moreover, editing associations with cancer stemness, DNA repair deficiency, and tumor immune infiltrations uncovered known and potential regulators of tumor features. We constructed an editing-mediated regulatory network, which revealed new editing modulators, including experimentally validated *TNRC6A*. Lastly, we demonstrated that EIs can act as biomarkers of both anti-PD1 immunotherapy and other cancer drugs, such as MEK and BRAF inhibitors. Collectively, iPEAPR enabled depicting RNA editing-dependent abnormal functions in cancer and can be applied to more diseases.

## Introduction

Millions of RNA editing sites, mainly adenosine-to-inosine (A-to-I) induced by adenosine deaminases acting on RNA (ADARs), have been identified in human [1–3]. Although most RNA editing sites (RES) are not evolutionarily conserved [4] and do not affect protein sequences [2], some functional A-to-I events have been reported in cancer, such as recoding events of *AZIN1* editing in liver or colorectal cancer [5, 6] and *RHOQ* editing in colorectal cancer [7], as well as *PKR* 3’UTR editing in lung cancer associated with metastasis [8]. Since the launch of The Cancer Genome Atlas (TCGA) project, several studies have identified clinically relevant A-to-I editing sites across cancer types by using TCGA RNA sequencing (RNA-seq) data [9–11]. Similarly, based on RNA-seq data of the Genotype-Tissue Expression (GTEx) project, new editing modulators have been identified [4]. Notably, RNA editing plays an important role in innate antiviral immune response and cancer stemness [12–14], which makes RNA editing related genes possible targets in cancer patients, but our knowledge is incomplete.

RNA editing enzymes, such as ADARs, are RNA binding proteins (RBPs), which have frequent interactions with each other [15, 16]. For example, hnRNPA2/B1 can enhance ADAR editing activity [17], HuR and ADARs cooperate to regulate RNA stability [18], and the DZF domain-containing protein ILF3 directly represses ADAR editing [19], among others [20]. RBPs interacting with ADARs may affect numerous cellular processes or pathways regulated by these RBPs. For example, ADAR may compete with m6A-related RBPs (e.g. METTL3, METTL14, and FTO) to modify adenosines [21]. Theoretically, editing enzymes may also interact with regulatory RNAs, such as enhancer RNAs (eRNAs) transcribed from enhancer elements or super enhancers (SE) [22]. Moreover, these interactions may occur in a cell-type specific manner [20], while most of above studies reports findings in a specific cell line.

However, there is a lack of systematic investigation of interactions between RNA editing and cancer hallmarks, functional gene sets, or functional genomic elements, such enhancers, RBP binding sites, and splicing regions, to further understand the roles of RNA editing in various cancer types. Recently, Roth et al. proposed the Alu editing index (AEI) to quantify the ADAR enzyme activity across the genome in each RNA-seq sample [23]. They found that AEI was mostly up-regulated in cancer [24]. But the AEI calculation mainly quantifies editing in Alu elements and requires raw sequencing data (large alignment BAM files) which hinders its application due to privacy and computing issues. We hypothesized that the AEI concept can be adapted to study interactions between RNA editing and, for example, RBPs and cancer hallmarks across tissues or cancer types, based on only site-level editing summary data, which are small in size for fast computation and without privacy issues. More importantly, studying these interactions may reveal novel important editing modulators.

Here, we developed iPEAPR (integrative pan-cancer/tissue editing activity of pathways and regions), a computational pipeline and web server to comprehensively profile and analyze RNA editing activities in functional genomic elements and pathways across cancers or tissues. Our analysis revealed highly heterogeneous editing activities across these features. Editing associations with cancer features, such as stemness, DNA damage deficiency, and immune infiltrations uncovered large numbers of significant and interesting regulators of pathophysiological functions in cancer. Moreover, we constructed a RBP regulatory network mediated by RNA editing, revealing known and novel biological functions supported by experimental evidence. Lastly, we demonstrated that EIs can act as biomarkers of both anti-PD1 immunotherapy and other cancer drugs, such as kinase inhibitors. iPEAPR can be further applied to other diseases, especially autoimmune diseases that are associated with abnormal ADAR activities [25, 26], and numerous pathways of interest.

## Results

### iPEAPR overview

We previously profiled RESs (97.4% A-to-I, **Fig. S1A**) in tumors and normal adjacent tissues (NAT) across 21 cancer types (**Fig. 1A**) from the TCGA project (https://gdc.cancer.gov<u>, see</u> **<u>Methods</u>**). About 31.4% (1,328,398 out of 4,234,680) of these RESs were specific to TCGA (termed cancerRES), compared to reported RESs from REDIportal (n= 4,668,508 ) [2], which was mostly based on cross-tissue healthy control samples from the GTEx project (https://gtexportal.org) (**Fig. S1B**) [4]. We further manually curated comprehensive annotations of the genome, transcriptome, and epigenome or epitranscriptome, including pathways or functional gene sets (such as cancer hallmarks), enhancer elements (including super enhancers), RBP binding sites (>200 RBPs), and m6A modification regions (**Fig. 1B**).

**Fig. 1.**
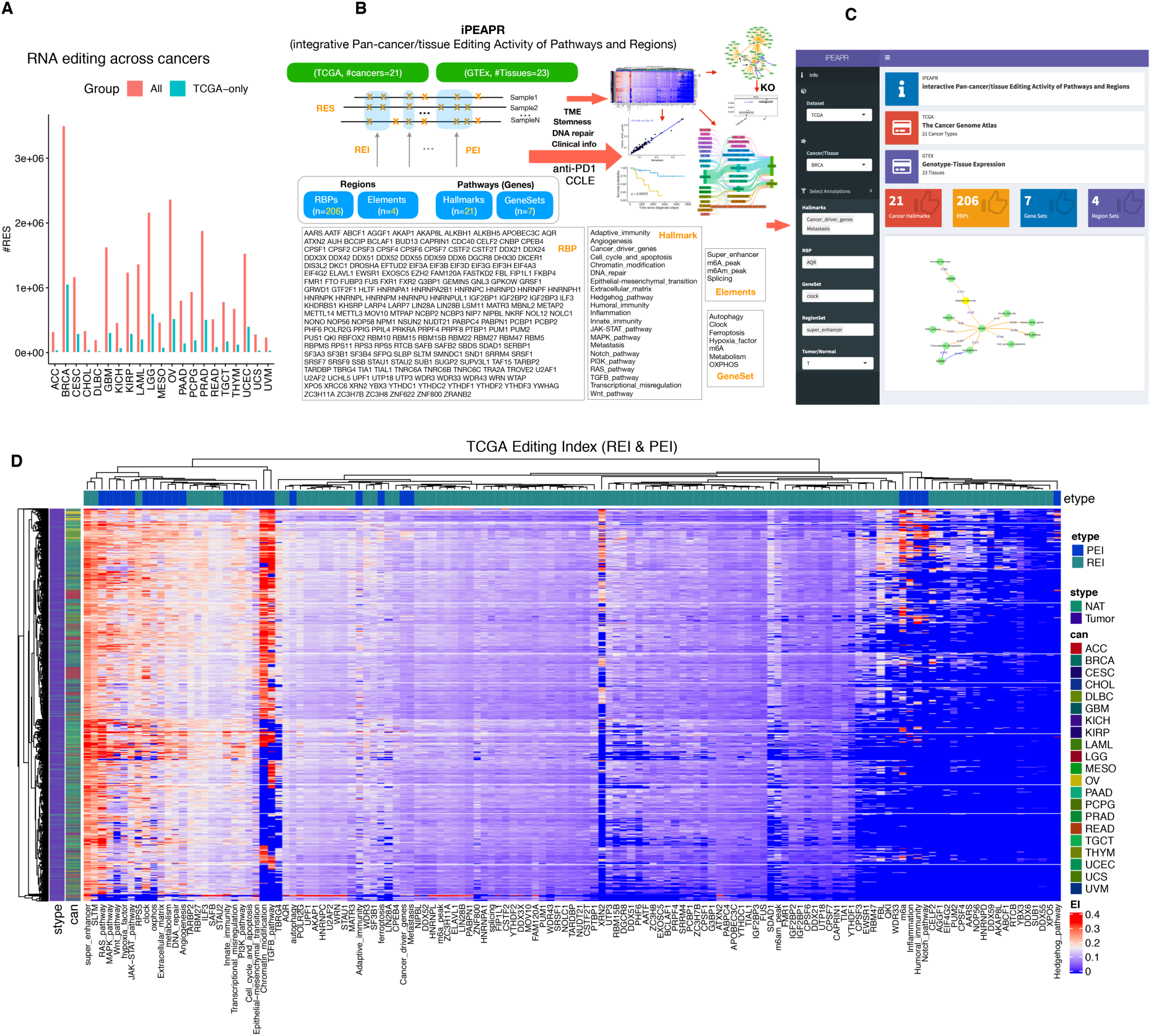
Overview of TCGA RNA editing and the iPEAPR framework. (**A**) Number of RESs (red color) or TCGA-only RESs (cancerRES, blue color) in each cancer. (**B**) Overview of the iPEAPR (integrative pan-cancer editing activity of pathways and regions) computational framework, based on 206 RBPs, four types of functional elements, 21 cancer hallmarks, and seven functional gene sets. Comprehensive analyses were conducted both for EI pairs and by combining EIs with sample features, including clinical information, TME, cancer stemness, and DNA repair deficiency. One newly identified modulator of RNA editing, TNRC6A, was experimentally validated by gene knockout. Furthermore, EIs as drug biomarkers were also identified for anti-PD1 and other oncologic or repurposed non-oncologic drugs. (**C**) The user-friendly and easy-to-use iPEAPR Portal for interactive analysis, built on all EIs calculated for TCGA and GTEx. (**D**) Overview of all REIs and PEIs (colored heatmap values) calculated in this study for each sample from 21 cancer types of TCGA. Super enhancers showed high REIs across cancer types. Rows are samples and columns are editing indexes (EIs). Cancer types and sample types were indicated on the left. RES: RNA editing site, EI: editing index, REI: regional editing index, PEI: pathway editing index, RBP: RNA binding protein, TME: tumor microenvironment, KO: knockout, ACC: Adrenocortical carcinoma, BRCA: Breast invasive carcinoma, CESC: Cervical squamous cell carcinoma and endocervical adenocarcinoma, CHOL: Cholangiocarcinoma, DLBC: Lymphoid Neoplasm Diffuse Large B-cell Lymphoma, GBM: Glioblastoma multiforme, KICH: Kidney Chromophobe, KIRP: Kidney renal papillary cell carcinoma, LAML: Acute Myeloid Leukemia, LGG: Brain Lower Grade Glioma, MESO: Mesothelioma, OV: Ovarian serous cystadenocarcinoma, PAAD: Pancreatic adenocarcinoma, PCPG: Pheochromocytoma and Paraganglioma, PRAD: Prostate adenocarcinoma, READ: Rectum adenocarcinoma, TGCT: Testicular Germ Cell Tumors, THYM: Thymoma, UCEC: Uterine Corpus Endometrial Carcinoma, UCS: Uterine Carcinosarcoma, UVM: Uveal Melanoma.

Based on the rich annotation information and millions of RESs from both TCGA and GTEx, we proposed a computational, integrative, and interactive analysis framework, iPEAPR (**Fig. 1B**, see **Methods**). We hypothesized that binding sites of different RBPs may have overall different RNA editing levels. Inspired by the Alu editing index (AEI) [23] to measure genome-wide ADAR enzyme activity by using raw read alignments (large BAM files) as input, we constructed a region editing index (REI), based on processed editing summary information (small text files), to measure editing enzyme activities in specific regions, for example, RBP binding sites, in each sample. Similarly, considering that pathways or gene sets may have members interacting with editing enzymes (also RBPs), we proposed a pathway editing index (PEI) to measure editing activity in each pathway or gene set. Then, we analyzed REI or PEI scores across cancer types from TCGA or normal tissues from the GTEx project. Finally, EI (REI or PEI) scores were combined with RBP expression levels to identify new editing modulators.

iPEAPR calculates REI for the following RBP-related annotations, including bound regions for each of 206 RBPs [15, 16], regions around m6A or m6Am (terminal m6A modification at the 5′ end of mRNA) modifications [27], splicing sites, and super enhancers. Enzymes of m6A modifications on RNA are an important group of RBPs. ADAR-induced A-to-I RNA editing can compete with m6A enzymes to modify adenosines [21]. RNA splicing sites are also bound by splicing factors, another important class of RBPs. Enhancers or super enhancers (SE) transcribe eRNAs (or seRNAs), which are bound by various RBPs and may act as scaffolds or guide molecules to perform long-range regulation of gene transcription [28]. In addition, iPEAPR calculates PEI for the following pathways or gene sets, including 21 cancer hallmarks [29], autophagy, circadian rhythm (clock genes), ferroptosis, hypoxia, m6A enzymes, metabolism, and OXPHOS (oxidative phosphorylation).

### iPEAPR Portal for interactive exploration of pan-cancer and cross-tissue EIs

To enable further exploration of our iPEAPR results and new analysis of our data, we built the iPEAPR Portal (**Fig. 1C**, freely available online at http://bioinfo-sysu.com:3838/sample-apps/ei), containing all EIs generated and analyzed in this work. In addition to those presented here as examples in this work, detailed analysis results for individual cancers or tissues are available on the iPEAPR Portal. Various types of interactive analyses and visualizations can be easily performed using the iPEAPR Portal to obtain more biological insight for future research.

### Differential but correlated RNA editing activities across different types of functional annotations

Overall, we observed heterogeneous editing activities across functional regions and gene sets in both tumors and normal samples from TCGA (**Fig. 1D**) and normal tissues from GTEx (**Fig. S1C**), suggesting considerably differential RNA editing across various annotations, which is consistent with our hypothesis. Particularly, we observed remarkably high EIs in the following regions or pathways, including super enhancers, SLTM (SAFB-Like Transcription Modulator), Chromatin_modification, RAS_pathway, and TGFB_pathway. Interestingly, cancerRES-based EIs were much smaller compared to EIs based on all RESs and were mainly observed in super enhancer, JAK-STAT, PI3K, Cell_cycle_and_apoptosis, m6A peaks, HNRNPC, and ELAVL1 (**Fig. S1D**).

On the other hand, positive pairwise EI correlations were observed in both TCGA (**Fig. 2A**, upper right triangle) and GTEx (**Fig. 2A**, lower left triangle), suggesting moderately coordinated RNA editing activities in functional regions across the genome, especially for RBPs, consistent with their cooperative RNA binding characteristics. Generally, cross-tissue pairwise EI correlations were weaker in TCGA cancers than in GTEx tissues (**Fig. 2B**). Robust pairwise EI correlations indicated that iPEAPR is able to estimate biologically meaningful EIs for different sets of functional regions.

**Fig. 2.**
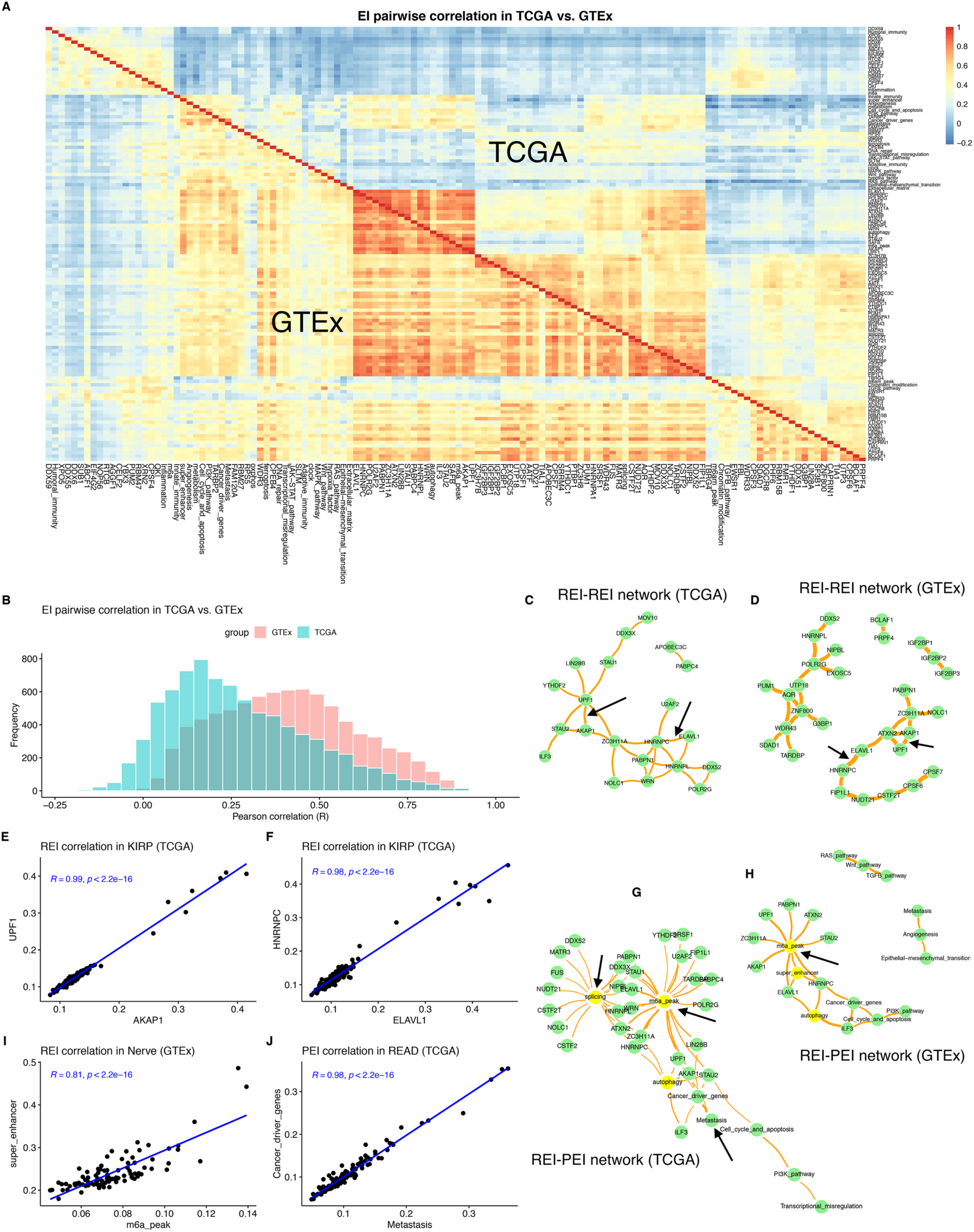
Pairwise EI correlations in TCGA cancers in comparison with GTEx tissues. (**A**) Heatmap of pairwise EI correlations in TCGA (upper right triangle), compared to GTEx (lower left triangle). A subset of RBPs showed the highest correlations. (**B**) Lower EI correlations in TCGA (green color) than in GTEx (red color), as also shown in (**A**). (**C-F**) The core subnetwork cross-cancer or cross-tissue significant REI correlations between RBPs in TCGA (**C**) and in GTEx (**D**). Green circles are RBPs, and orange edges are the scaled sum of correlation coefficients (R) across different cancer types of the same REI pair. Black arrows are pairs described in main text and with almost perfect correlations, including UPF1-AKAP1 (**E**) and HNRNPC-ELAVL1 (HuR) (**F**). (**G-J**) The core subnetwork significant REI-PEI or PEI-PEI correlations (except those shown in **C** and **D**) in TCGA (**G**) and in GTEx (**H**). Circles are RBPs or cancer hallmarks, yellow circles are selected functional set of elements or genes, and orange edges are the scaled sum of correlation coefficients (R) across different cancer types of the same EI pair. Black arrows are pairs described in main text, including m6A_peak-super_enhancer (**I**) and Metastasis-Cancer_driver_genes (**J**).

### Editing index correlation network analysis revealed new biological functions

To uncover possible new functional RBP clusters, we calculated pairwise EI correlations of RBPs in each cancer (TCGA) or tissue (GTEx). We then visualized the network (editing index correlation network, or EICN) of those RBP pairs with strong correlation (R-squared >= 0.6) in at least 20 cancer types (TCGA, **Fig. 2C**) or tissues (GTEx, **Fig. 2D**). We identified both known and novel functional clusters of RBPs.

Notably, we observed almost perfect correlations between AKAP1 and UPF1 (**Fig. 2E**), and between ELAVL1 (or HuR) and HNRNPC (**Fig. 2F**), across all cancer types, suggesting close relationship between both RBP pairs. This correlation was also observed across normal tissues (**Fig. 2D**). Significant overlapping of binding sites by ELAVL1 and HNRNPC were observed in a previous report [30]. However, the specific relationship between AKAP1 and UPF1 is unknown. UPF1 functions as a component of the RNA surveillance complex of mRNA-mediated decay, and AKAP1 binds to protein kinase A to anchor it to specific cellular locations. We noticed that both AKAP1 and UPF1 show physical interactions with PABPC1, but only UPF1 has physical interaction with the editing enzyme ADAR (the STRING database, https://string-db.org). Moreover, both UPF1 and AKAP1 were indicated in binding RNA guanine quadruplexes (G4) [31]. Therefore, AKAP1 and UPF1 may cooperate with ADAR to bind G4 structures and edit adenosines in the G4 loops to resolve G4s.

Similarly, we built EICN network between RBPs and functional sets, or only within functional sets (**Fig. 2G**). In cancer, the m6A_peak showed associations with various RBPs, including UPF1, WRN, STAU1, STAU2, and PABPN1, many of which were also identified across normal tissues (**Fig. 2H**). In contrast, most associations of splicing REIs were lost in normal tissues. Moreover, in GTEx, we observed strong correlation between m6A_peak and super enhancers (one example shown in **Fig. 2I**), consistent with recent findings that abundant m6As on seRNAs regulate expression levels of seRNAs and their target genes [32]. Together with high REIs we found in enhancers, our observations indicate that RNA editing probably regulates eRNA abundance by interfering m6A modifications. Additionally, in TCGA, we observed a strong correlation between two cancer hallmarks, Metastasis and Cancer_driver_genes (note that only a small number of overlapping genes; one correlation example shown in **Fig. 2J**), correlation of which was also found in normal tissues.

### Prognostic EIs in multiple cancer types

When compared to adjacent normal tissues, tumor EI scores were generally elevated in BRCA, but decreased in GBM, CHOL, and KICH (**Fig. S2**), which is different from previous results based on genome-wide editing activities that showed elevated AEIs in most cancer types [24]. Notably, angiogenesis PEI was increased in five cancer types, and Epithelial-mesenchymal_transition (EMT) PEI was consistently decreased in three cancer types, including BRCA, KIRP, and CHOL.

We next tried to survey prognostic EIs across cancers, because EIs were aggregated RNA editing features, which largely reduced the burden of multiple correction of significance for genome-wide RNA editing analysis. For EIs based on all RESs, a total of 12 hallmarks, gene sets, or RBPs were significantly associated with patient survival (FDR<0.1, **Fig. 3A**). The most significant one was EI of EMT in BRCA (*P*=5.1e-6, **Fig. 3B**; low EI showed worse outcome, concordant with consistently lower EMT EIs in tumor samples in comparison with normal controls), followed by EIs of Wnt_pathway, splicing, NUDT21, and MAPK_pathway. EMT associated editing was associated with multiple cancer types [8]. Different hallmarks were prognostic in different cancers, possibly reflecting their cancer type-specific roles. Additionally, we found that four cancerRES-based EIs were significantly associated with patient survival (FDR<0.1, **Fig. 3C**). The most significant EI for prognosis was still EMT in BRCA (*P*=2.5e-5), followed by EIs of super_enhancer in ACC (*P*= 3.7e-4, **Fig. 3D**).

**Fig. 3.**
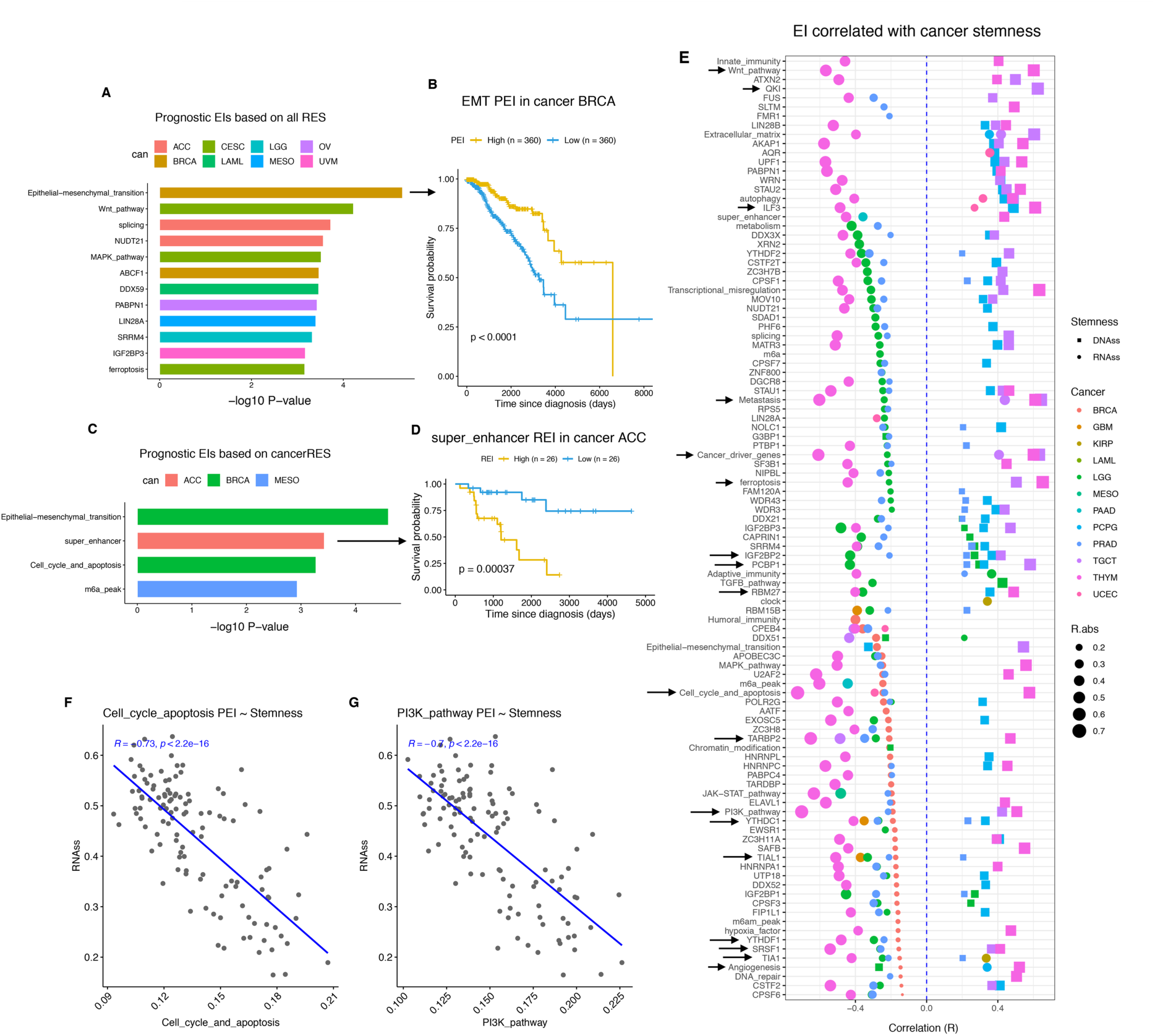
Prognostic EIs and EIs associated with cancer cell stemness. (**A-B**) Significant EIs that were prognostic in different cancer types, based on all RESs (**A**), and Kaplan-Meier plot for epithelial-mesenchymal transition (EMT) in BRCA (**B**). (**C-D**) Significant EIs that were prognostic in different cancer types, based on cancerRESs only (**C**), and Kaplan-Meier plot for super_enhancer in ACC (**D**). (**E-G**) Significant correlations between EIs and cancer stemness across cancer types (**E**). Black arrows indicate features with high EI-stemness correlations described in the text, including (**F**) Cell_cycle_and_apoptosis-RNAss and PI3K_pathway-RNAss (**G**). Symbol shapes are stemness types. Colors are cancer types. Symbol sizes are absolute correlation coefficients (R). RNAss: RNA expression-based stemness score. DNAss: DNA methylation-based stemness score.

### Stemness-associated RNA editing activities reveal new players in cancer stem cells

RNA editing contributes to cancer stemness [12, 14] and is associated with cancer metastasis via EMT [8]. Therefore, we investigated the EIs associated with cancer stemness reported by a previous study across >30 cancer types [33]. Surprisingly, we found that RNA expression-based stemness scores (RNAss) were negatively associated with EIs, while DNA methylation-based stemness scores (DNAss) were positively associated with EIs (**Fig. 3E**). Our RNA editing index may better explain the observed discrepancies between these two types of stemness in the original study [33].

In total, we observed 378 pairs (247 EI-RNAss and 131 EI-DNAss, **Table S1**) with significant (FDR <0.01) correlations between EIs and stemness scores, mainly in THYM (100), LGG (72), PRAD (72), BRCA (42), PCPG (40), and TGCT (37). Interestingly, four of the five negative EI-DNAss pairs, such as Angiogenesis, were in LGG, and the remaining one (EMT) in PCPG (paraganglioma). The top three positive EI-RNAss pairs, such as Metastasis-RNAss, were from TGCT. For EI-RNAss partners, YTHDC1 (m6A reader), TIA1, TIAL1, TARBP2, and CPEB4 were all observed in five cancer types, in addition to YTHDF1 (m6A reader) and SRSF1 that appeared in four cancer types. For EI-DNAss pairs, PCBP1 and IGF2BP2 were both observed in four cancer types. Interestingly, the uncharacterized RBP, RBM27, showed significant associations with stemness in four cancer types (THYM, LAML, LGG, and PCPG), which probably indicates its critical function in stemness regulation.

The most significant stemness-associated EIs were two cancer hallmarks, Cell_cycle_and_apoptosis (R=-0.725, *P*=1.1e-20; **Fig. 3F**) and PI3K_pathway (R=-0.702, *P*=5.3e-19; **Fig. 3G**), both showing negative correlations with RNAss stemness in THYM. EIs of ferroptosis, Metastatsis, and Cancer_driver_genes were positively associated with DNAss stemness. Interestingly, QKI, U2AF2, and ILF3 were among the top 20 best correlation EI partners with stemness. QKI [34] and U2AF2 [35] are known to regulate stemness and angiogenesis of cancer stem cells, respectively. ILF3 regulates the EIF2AK2-encoded PKR abundance to induce EMT in a editing-dependent manner [8].

### Tumor immune infiltration associated RNA editing activities reveal new players in TME

RNA editing plays an important role in innate antiviral immune response [13, 14]. Therefore, we further examined the association between EI and immune infiltration in tumor microenvironment (TME) (**Fig. 4A**, **Table S2**). Remarkably, we observed negative EI-TME correlations (FDR <0.01) predominantly for the cancer hallmark Adaptive_immunity across cancer types, with the most significant correlation with Neutrophils in UCEC (R= -0.55, *P*=6.2e-31; **Fig. 4B**). In addition, EI-TME correlations for the uncharacterized RNA helicase DDX51 were mostly negative and seven of the nine pairs were observed in LGG, indicating the important immune-related role of DDX51 in LGG.

**Fig. 4.**
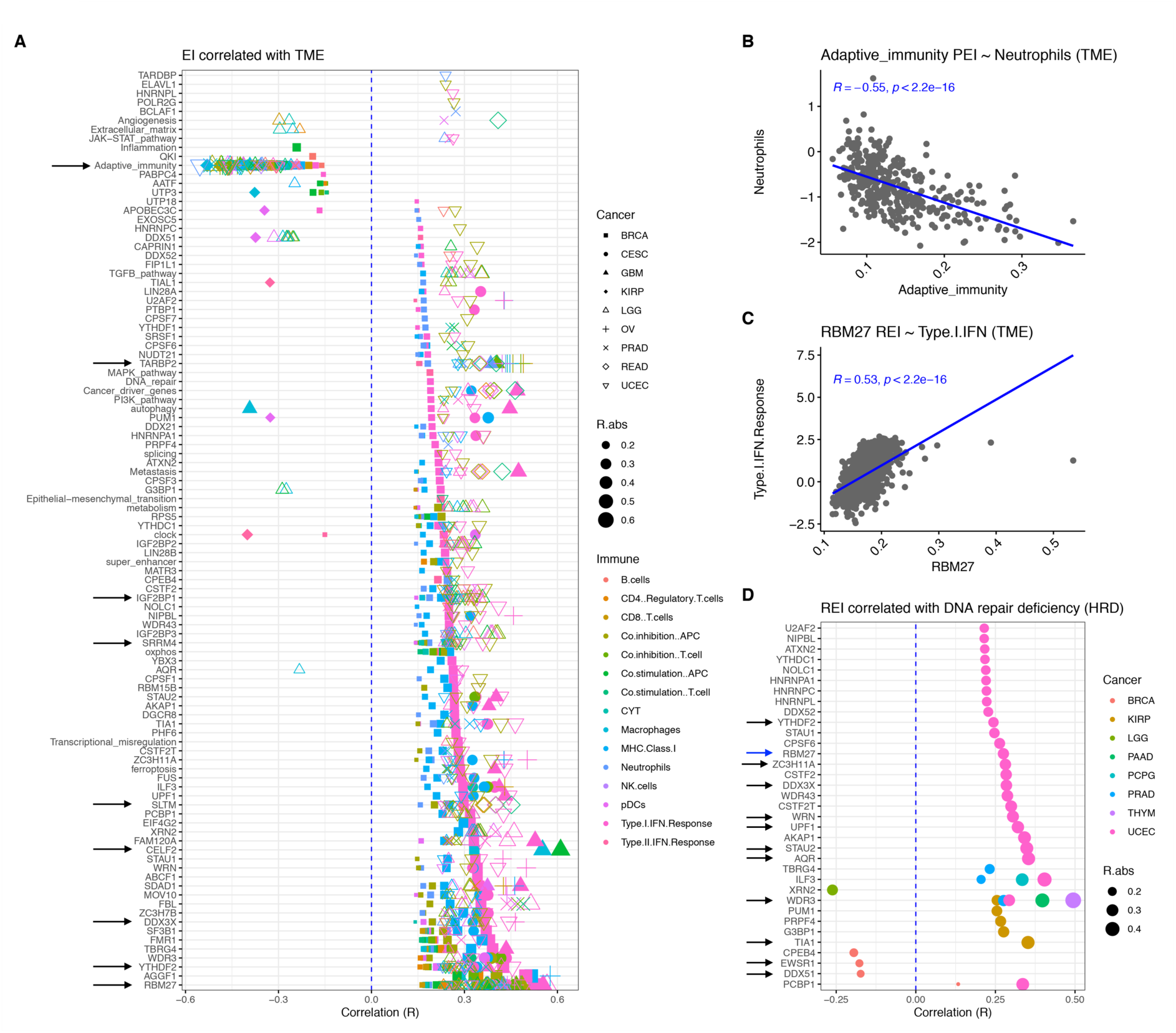
EIs associated with infiltrated immune cells in tumor microenvironment (TME) and DNA repair deficiency (HRD) in tumors. (**A-C**) Significant correlations between EIs and fractions of infiltrated immune cells in TMEs across cancer types (**A**). Symbol shape are cancer types. Colors are TME features. Symbol sizes are absolute correlation coefficients. Two examples were shown, Adaptive_immunity PEI with Neutrophils (**B**) and RBM27 REI with Type.I.IFN.Response (**C**). (**D**) Significant correlations between EIs and HRD scores across cancer types. Symbol sizes are absolute correlation coefficients. Colors are cancer types. Symbol sizes are absolute correlation coefficients.

In contrast, most EI-TME associations were positive and the highest correlation was from CELF2 with co-stimulation of APCs (antigen-presenting cells) in GBM (R=0.610, *P*=2.3e-7). CELF2 is induced in stimulated T cells and regulates alternative polyadenylation (APA) in immune responses [36]. Surprisingly, the uncharacterized RBM27 was the most frequent partner of EI-TME pairs, including MHC class I in BRCA and co-inhibition of T cells in LGG, followed by YTHDF2 (m6A reader, n= 37), IGF2BP1 (n=28), TARBP2 (n=23), DDX3X (n=20), SLTM (n=20), and SRRM4 (n=20). The most significant correlation was from RBM27 with Type I IFN (interferon) response in BRCA (R= 0.53, *P*=1.7e-74; **Fig. 4C**). Of note, YTHDF2 regulates NK cell-mediated antitumor and antiviral immunity in mouse [37], while DDX3X regulate type I interferon response and cytotoxic T cells and dendritic cells infiltration via MDA5 and ADAR1 in breast cancer [38]. Immune-related roles in TME for other RBPs, especially RBM27, mentioned above warrant further research.

### RNA editing-dependent DNA repair by WDR3 across cancer types

DNA repair requires ADAR-mediated RNA editing in DNA:RNA hybrids (R-loop) [39]. To discover new RNA editing-dependent regulators of DNA repair, we examined the associations between RBP EIs with homologous recombination deficiency (HRD) scores across cancer types from TCGA. We identified 23 significant (FDR <0.01) REI-HRD pairs, most of which only observed in UCEC, including the uncharacterized RBM27 (**Fig. 4D**). Notably, about 50% of these RBPs are known to function during DNA damage repair, suggesting the reliability of our results. In addition to several other identified RBPs, such as YTHDF2, DDX51, DDX3X, EWSR1, TIA1, STAU2, and ZC3H11A (see **Supplementary Information** for related publications), WRN is well-known and critical for DNA repair [40]; deficiency of the RNA helicase AQR induces R-loop accumulation which results in DNA damage [41]; UPF1 promotes R-loops to enhance DNA break repair [42].

Remarkably, we noticed that WDR3 showed significant positive correlations in four cancer types (THYM, PAAD, UCEC, and PRAD), with suggestive correlations in most other cancer types (**Fig. S3**). The WDR3 was indicated to be possibly involved in DNA repair because it binds regions near strand breaks after multiple types of DNA damages, yet with unknown mechanism [43]. Our findings suggest that WDR3-related DNA repair is probably RNA editing-dependent.

### Integrating gene expression in editing-mediated regulatory network reveals new modulators of RNA editing

To further investigate editing-mediated interactions with RBPs, we examined pairwise correlation between EIs and RBP expression levels (EI-RBP expression pairs, see **Methods**). A total of 8,172 and 17,759 significant EI-RBP pairs (|R|>=0.3, FDR<0.05; identified in at least two cancers or tissues) were identified in 20 cancer types of TCGA (**Fig. 5A**) and in 21 normal tissues of GTEx (**Fig. 5B**), respectively. For TCGA, the largest source of EI-RBP pairs was THYM and the smallest was UCEC. Interestingly, more than 70% EI-RBP pairs of brain cancers (both LGG and GBM) showed negative correlations. In contrast to TCGA, for GTEx, many tissues showed more negative than positive correlations, and the brain and colon showed large percentage of positive correlations.

**Fig. 5.**
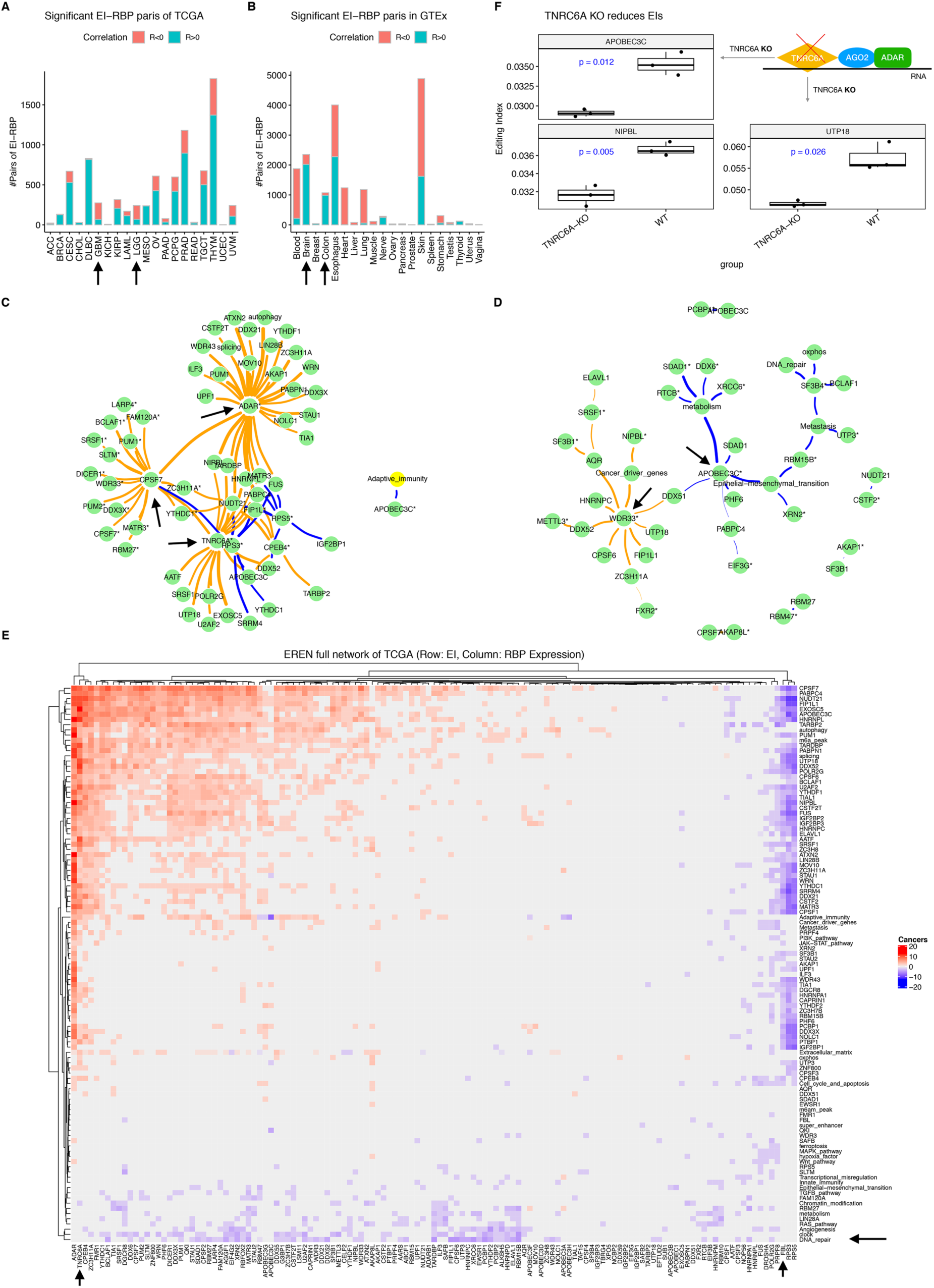
EI-RBP regulatory network (EREN) reveals new RNA editing modulators supported by experimental evidence. (**A-B**) Frequencies of all significant EI-RBP pairs in each TCGA cancer type (**A**) or each GTEx tissue (**B**). (**C-D**) The core subnetwork of EREN, with each EI-RBP pair showing in at least eight cancer types in TCGA (**C**) or tissues in GTEx (**D**). RBP expression nodes are marked with asterisks. Edges are scaled sum of correlation coefficients (R). Orange means positive correlation and blue means negative correlation. Black arrows indicate outstanding features described in the main text, including ADAR, CPSF7, and TNRC6A in TCGA and WDR33 and APOBEC3C in GTEx. (**E**) The full EREN network of TCGA showed as a heatmap. Rows are EIs features and columns are RBP expression features. Colors represent cancer frequencies of each significant EI-RBP pair across TCGA cancer types. Red stands for positive association and blue stands for negative association. Black arrows indicate outstanding features described in the main text. (**F**) TNRC6A knockout (KO) reduced EIs of TNRC6A-associated RBPs, such as APOBEC3C, NIPBL, and UTP18, in the TCGA EREN core subnetwork shown in (**C**).

For TCGA, as expected, ADAR was among the most frequent partners of this editing-mediated RBP regulatory network (EREN, a sub-network was shown in **Fig. 5C**). Among the five positive EI-RBP pairs appeared in 12 cancer types, four involved ADAR (PUM1_ADAR, NIPBL_ADAR, HNRNPL_ADAR, and ATXN2_ADAR), and the remaining one was EXOSC5_TNRC6A, which warrants further research. TNRC6A functions in post-transcriptional silencing by miRNA or RNA interference [44] while EXOSC5 encodes a subunit of the RNA exosome complex. Interestingly, among the six negative EI-RBP pairs appeared in at least nine cancer types, either RPS3 or RPS5 was involved (APOBEC3C_RPS3, FIP1L1_RPS3, FIP1L1_RPS5, SRRM4_RPS3, NUDT21_RPS3, and NUDT21_RPS5), which suggests their roles in regulating target RBPs mediated by RNA editing.

Notably, REI of CPSF7 was associated with expression of plenty of RBPs, including CPSF7 itself, RBM27, DICER1, SLTM, and SRSF1, and in contrast to TNRC6A, whose expression potentially regulated large numbers of RBP REIs (**Fig. 5C**). CPSF7 is required for the pre-mRNA 3’-processing, i.e. cleavage and polyadenylation [45]. In addition to regulation of REIs of various RBPs and splicing sites, ADAR was also associated with autophagy PEI, suggesting cross-talk between RNA editing and autophagy. Moreover, APOBEC3C (apolipoprotein B mRNA editing enzyme catalytic subunit 3C) potentially regulates adaptive immunity PEI.

For GTEx, ADAR was not frequently associated with specific EIs across tissues (**Fig. 5D**), although many ADAR-involved EI-RBP pairs were identified (**Fig. S4**). Interestingly, WDR33 expression was positive correlated with EIs of Cancer_driver_genes and many RBPs, including AQR, HNRNPC, DDX52, DDX51, CPSF6, and FIP1L1, while expression levels of APOBEC3C and SF3B4 were negatively associated with EIs of many RBPs, such as DDX51 and PABPC4, cancer hallmarks, including EMT, DNA_repair, and Metastasis, and metabolism genes. APOBEC3C REI was also correlated with PCBP1 expression.

Besides, the full EREN network revealed more biology of EI-RBP interactions (**Fig. 5E**). FBL and PRPF8 were very similar to RPS3 and RPS5, negatively associated with large number of REIs of RBPs. In contrast, TNRC6A, CPEB4, ZC3H11A, FMR1, and YTHDC1 (m6A reader) were similar to ADAR, positively associated with large number of both REIs and PEIs. Remarkably, PEIs of several cancer hallmarks, including DNA repair, angiogenesis, RAS pathway, together with functional gene sets, such as clock and metabolism, consistently showed negative correlation with expression of various RBPs. We believe further in-depth of this EREN network should shed light on more biological processes related to RNA editing activity.

### TNRC6A knockout reduces REIs of various RBPs identified in the pan-cancer EREN network

To experimentally validate our findings in the pan-cancer EREN network, we examined the EI impact of TNRC6A (trinucleotide repeat containing adaptor 6A), which regulates pan-cancer REIs of 16 RBPs, such as APOBEC3C, EXOSC5, DDX52, HNRNPL, CPSF7, SRSF1, U2AF2, and TARBP2. Of note, several of these RBPs, such as SRSF1, U2AF2, and TARBP2, were associated with cancer stemness as shown above. Consistent with the observed similarity between TNRC6A and ADAR (**Fig. 5E**), TNRC6A works with argonaute protein 2 (AGO2), which interacts with ADAR [46], to degrade mRNAs via short interfering RNAs (siRNAs) [44]. We compared the REIs of these RBPs before and after TNRC6A knockout in HCT-116 colon cancer cells [47]. As predicted, REIs of all TNRC6A-associated RBPs, except MATR3, in the EREN network were reduced (**Fig. 5F**, **Fig. S5**).

### REIs predict anti-PD1 responses and treatment effects of other cancer drugs and suggest possible drug mechanisms

Immunotherapies by immune checkpoint inhibitors (ICIs), such as anti-PD1, are very promising cancer treatment strategies yet with a diverse range of patient responses and few effective and reliable biomarkers [48]. To demonstrate potential clinical usage of iPEAPR, we investigated the relationship between EI and patient treatment response of anti-PD1. Then, we calculated EIs for melanoma patients before and during anti-PD1 therapy [49]. Interestingly, we observed significant correlation between anti-PD1 response (ordered as progressive disease [PD], stable disease [SD], and partial response or complete response [PRCR]) during treatment and EIs of Adaptive_immunity (negative correlation, *P*=0.0036), Cell_cycle_and_apoptosis (positive, *P*=0.053), Metabolism (positive, *P*=0.04), OXPHOS (positive, *P*=0.0065), FUBP3 (positive, *P*=0.0052), and PCPB2 (negative, *P*=0.0022) (**Fig. 6A**). PEIs of Adaptive_immunity, Cell_cycle_and_apoptosis, Metabolism, or OXPHOS possibly affected genes included by them, while REIs of FUBP3 or PCBP2 possibly affected their target RNA binding affinity. However, the same correlations were not significant, although showing similar trends, for patients before treatment (**Fig. S6A**). Interestingly, increased FUBP3 protein level was observed after blockade of PD1-PD-L1 interaction [50]. Currently, the mechanisms behind the link between anti-PD1 and FUBP3 or PCBP2 are unknown and require further experimental work.

**Fig. 6.**
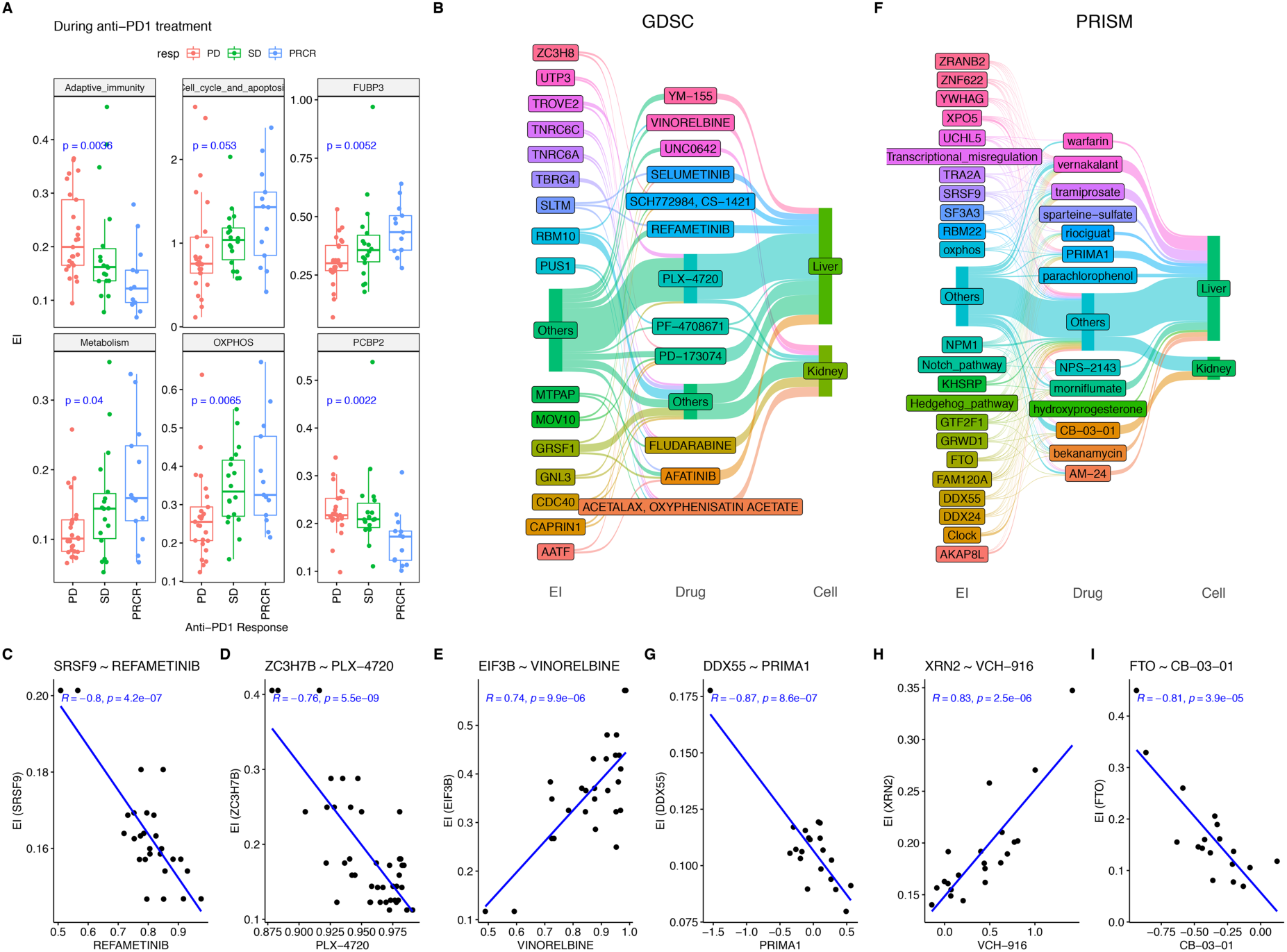
EIs are biomarkers of anti-PD1 immunotherapy and other cancer drugs. (**A**) During anti-PD1 treatment, six types of EIs were associated with patient responses, including Adaptive_immunity, OXPHOS, and FUBP3. PD, progressive disease; SD, stable disease; PRCR, partial response or complete response. (**B-E**) In the GDSC drug screening resource, EIs (the first column) were associated with various types of cancer drug treatment effects (the second column) in liver or kidney cancer cells (the third column), such as (**C**) SRSF9 with REFAMENTINIB (a MEK inbhitor), (**D**) ZC3H7B with PLX-4720 (a BRAF inhibitor), and (**E**) EIF3B with VINORELBINE (a chemotherapy drug). EIs or drugs with only one link were grouped into ‘Others’. (**F-I**) Similar to (**B**), in the PRISM drug repurposing screening resource, EIs were associated with various types of repurposed non-oncologic drug treatment effects in liver or kidney cancer cells, such as (**G**) DDX55 with PRIMA1 (TP53 inhibitor), (**H**) XRN2 with VCH-916 (HCV inhibitor), and (**I**) FTO with CB-03-01 (androgen receptor antagonist). EIs or drugs with <5 links were grouped into ‘Others’.

We further examined EI relationships with treatment of other drugs, including mostly oncological and repurposed non-oncological drugs from GDSC [51] and PRISM [52] resources, respectively, in the liver and kidney cell lines from the Cancer Cell Line Encyclopedia (CCLE) [53]. For drug treatments from GDSC, we identified 59 and 26 pairs of significant correlation between EIs and responses of drugs in liver and kidney, respectively (**Fig. 6B**). For example, in liver cells, the response of REFAMETINIB (a MEK inhibitor) was highly correlated with REI of SRSF9 (**Fig. 6C**), and PLX-4720 (a BRAF inhibitor) response was correlated with REI of ZC3H7B (**Fig. 6D**). In kidney cells, VINORELBINE (a chemotherapy drug) response was positively correlated with REI of EIF3B (**Fig. 6E**), among others. Only the correlation between PLX-4720 response and REI of TROVE2 (Ro60 autoantigen) was shared between liver and kidney cells, indicating tissue-specific drug effects and EIs. On the other hand, we identified 274 and 60 pairs of significant correlation between EIs and responses of drugs in liver and kidney, respectively (**Fig. 6F**). For example, PRIMA1 (TP53 inhibitor) response in liver was correlated with REI of DDX55 (**Fig. 6G**) and VCH-916 (HCV inhibitor) response was correlated with REI of XRN2 (**Fig. 6H**). In kidney cells, CB-03-01 (androgen receptor antagonist) response was correlated with REI of FTO (**Fig. 6I**).

## Discussion

In this work, we developed a computational and interactive framework, iPAEPR, to study the interactions between RNA editing and cancer hallmarks, RBPs, enhancers, and other functional elements. Studying associations of editing activities of these features with various tumor sample features across cancer types enabled us to pinpoint possible abnormal functions in each cancer, and to identify new regulators of both RNA editing and tumor features, including stemness, DNA repair, and immune infiltration in TME. Moreover, we demonstrated that EIs can act as biomarkers of both anti-PD1 immunotherapy and other cancer drugs, such as kinase inhibitors, and may help reveal possible mechanisms behind drugs. Besides cancer, many other diseases were characterized by aberrant ADAR activities, such as autoimmune diseases [25, 26], and may benefit from iPEAPR analysis.

We were surprised that REIs of super enhancer regions were among the highest across cancer types, compared to other features. The reason behind this observation is unclear. It is possible that ADAR may have unknown connections with enhancer RNAs via interactions with specific RBPs, such as m6A readers [32]. Since the FANTOM (Functional ANnoTation Of the Mammalian genome) project [22] produced a large volume of CAGE (cap analysis gene expression) sequencing data, which were specifically used to detect enhancers and may be utilized to further explore enhancer RNA editing to uncover their connections.

Compared to previous cancer studies based on only RESs from the REDIportal or the RADAR database [9–11], we found that cancerRES (found only in TCGA) added values to RES analysis in cancer. For example, cancerRES-based prognosis analysis revealed that super_enhancer and m6A_peak REIs were able to predict patient outcomes.

Our EREN network analysis revealed large numbers of potential interactions within about 200 RBPs and also between RBP and cancer hallmarks and functional elements, providing interesting directions for future research. EREN can be further expanded to include >1500 RBPs [15] and uncover their currently unknown functions, especially for most of the uncharacterized RBPs. More importantly, many of EI associations with tumor stemness, DNA repair, and TME features were not reported previously. Based on the large number of known associations identified by our approaches, the remaining EI associations should provide useful clues for future experimental designs and validations.

Several limitations represent possible further improvement of this analysis. First, PEI or REI may not be accurate estimations of ADAR activity in pathways or functional gene sets with incomplete annotations. For example, the current list of m6A enzymes or related genes have been actively being updated. Second, expanding the EREN network analysis to all gene expression, instead of only RBP gene expression, may uncover more RNA editing regulators and new findings. Moreover, new pathway and gene set annotations can be integrated into the analysis workflow, such as more gene sets or pathways from MSigDB (https://www.gsea-msigdb.org/gsea/msigdb/), KEGG (https://www.genome.jp/kegg/), or Reactome (https://reactome.org/). Lastly, besides the experimentally supported TNRC6A in regulating RNA editing, many other proposed novel functions of RNA editing or RNA binding proteins need further verification by wetlab experiments. For example, we propose that WDR3 probably regulates DNA repair via ADAR-mediated RNA editing in the R-loop structures of DNA break sites [39].

Collectively, we believe that iPEAPR can be further explored interactively for all cancer types to obtain biological insight to guide experimental designs, and it should be able to applied to more diseases to pinpoint possible pathogenic factors responsible for patient manifestations.

## Methods

### Data collection

TCGA RNA-seq raw reads, germline and somatic mutations, and sample clinical information were download from the GDC Portal (https://portal.gdc.cancer.gov). TCGA gene expression levels in TPM, stemness scores, and DNA repair deficiency scores were downloaded from the Xena database (https://xenabrowser.net). GTEx samples clinical information was obtained from dbGAP (https://dbgap.ncbi.nlm.nih.gov) with accession number phs000424.v6. RNA editing sites of GTEx were obtained from the REDIportal database [2]. RNA-seq raw reads for TNRC6A wildtype and knockout samples (n=6) were obtained from SRA (accession number: PRJNA682956). RNA-seq for anti-PD1 treated melanoma samples (n=109) were downloaded from SRP094781. RNA-seq for CCLE samples were downloaded from PRJNA523380. GDSC and PRISM drug screen effects were downloaded from DepMap (https://depmap.org).

### TCGA RNA editing sites identification and quality control

TCGA RNA editing sites in each cancer were identified previously [54]. Briefly, SPRINT [55] was applied to each sample to identify possible RESs, using hg38 as the reference and default parameters. Initial RESs were filtered as following: (i) detected in at least three samples in each cancer; (ii) detected in at least two cancer types; (iii) not overlapped with germline mutations; (iv) not overlapped with somatic mutations. Then, remaining RESs (N=4,234,680 unique sites) were merged and annotated to genes and dbSNP150 by ANNOVAR [56]. RESs overlapped with SNPs with allele frequencies >= 0.01 were excluded. RESs not found in the REDIportal database [2] were defined as TCGA cancer specific RESs.

### Curation of cancer hallmarks and functional gene sets

A total of 21 cancer hallmarks with different number of annotated genes were obtained from [29]. In addition, seven functional gene sets were manually curated from the following studies: autophagy [57], circadian rhythm [58], ferroptosis [59], hypoxia-induced factors [60], m6A enzymes and related genes [61], metabolism [62], and OXPHOS [62].

### Curation of RBP sites, m6A sites, splicing sites, enhancers and super enhancers

Binding sites for a total of 206 RBPs were obtained from the ENCODE project [15] and the POSTAR2 database [16]. Peaks of m6A and m6Am modifications across various tissues were obtained from [27]. Splicing sites were defined as regions centered on exon boundaries (GENCODE v28, https://www.gencodegenes.org) with 3-bp flanking each side. A subset of FANTOM-derived [22] super enhancers in cancer were obtained from [63].

### iPEAPR framework and the web Portal construction

First, to calculate PEI, genes annotated to each cancer hallmark or functional gene set were intersected with genes annotated with RESs to obtain overlapped RESs in each sample. To calculate REI, genomic regions belonging to each set of functional elements were intersected with positions of RESs in each sample directly to obtain overlapped RESs. Then, PEI or REI was estimated by dividing total edited bases by total sequenced bases of those overlapped RESs. For each sample, to obtain relatively robust PEI or REI estimation, a minimum of 50 RESs were required for each PEI or REI calculation. To speed up the calculation for a large number of cancer types and samples, iPEAPR adopted a parallel computational mode. The iPEAPR Portal was built by using R shiny (https://www.rstudio.com/products/shiny/shiny-server) and was based on all EIs calculated by iPEAPR in this study.

### Pairwise EI correlation analysis in TCGA and GTEx

Pairwise EI correlations (Pearson’s R) between REIs, REI-PEI, or PEI-PEI were calculated in each cancer type (TCGA) or each tissue (GTEx). Since many pairwise EIs showed high correlations, strict criteria of correlation were set for EIs to construct further network visualization. For all RES-based EI correlation network visualization, top correlations were selected by requiring FDR<1e-5, absolute R>=0.6, and identified in at least 20 cancer types or tissues. For cancerRES-based EI network visualization, top correlations were selected by requiring FDR<1e-5, absolute R>=0.5, and identified in at least 10 cancer types or tissues.

### Association analysis between EI and cancer stemness, HRD, or TME

Pearson correlations between EIs and tumor sample features (cancer stemness, HRD, or TME) were calculated in each cancer type. Significant EI-feature pairs with FDR<0.01 across cancer types were used for visualization and downstream analysis.

### Editing-mediated RBP regulator network (EREN) analysis

First, TCGA expression levels of all RBPs collected for REI, together with ADARs and APOBECs, were extracted from the downloaded full expression matrix. Then, pairwise Pearson correlation analysis was performed between each EI and the expression of each RBP (EI-RBP expression pair). Significant correlated EI-RBP expression pairs were defined as Pearson correlation coefficient R>=0.3 and FDR<0.05. We further required EI-RBP expression pairs appeared in at least two cancer types. For core subnetwork visualization, only those pairs appeared in more than eight cancer types were selected.

### RBP-based RNA-editing index analysis for TNRC6A wildtype and knockout samples

RNA-seq reads were aligned to the hg38 reference genome by HISAT2 (http://daehwankimlab.github.io/hisat2, v2.2.1). READ alignments were then used to produce base pileup profiles by using samtools (v1.9). The max RNA-seq read depth was set to 10,000. Polymorphic positions from dbSNP150 with minor allele frequency >=10% were excluded before intersecting pileup profiles with RBP binding regions for REI calculation. To calculate REIs for TNRC6A samples, RNA editing sites with a minimum editing ratio of 0.005 were used.

### EIs as biomarkers of drug treatment in patients and cell lines

For melanoma patients treated with anti-PD1 therapy, we compared EIs among different groups of patient responses, including progressive disease, stable disease, or partial response or complete response. For drug treatments in CCLE cell lines, we correlated EIs with drug effects in AUC values of response curves or log fold changes before and after treatments. *P*-values were adjusted by controlling FDR.

### Survival analysis, other bioinformatics analysis, and data visualization

Differential EIs were obtained by comparing EIs of tumors with NATs using *t*-test in R language. Survival analysis was performed and visualized by the R package TCGAbiolinks [64], by comparing samples with EI higher or lower than the 0.67 or 0.33 EI quantile across samples. *P*-values were adjusted by R function p.adjust by false discovery control (FDR). Heatmap data visualizations were visualized by R package ComplextHeatmap (v2.2) or pheatmap (v1.0). Drug biomarkers were visualized by ggsankey (v0.0.99). Networks were visualized by R package igraph (v1.2.6). Other visualizations were performed by R package ggpubr (v0.4.0).

## Data availability

All data generated or analyzed during this study are included in this published article and its supplementary information files. The iPEAPR portal can be accessed at http://bioinfo-sysu.com:3838/sample-apps/ei.

## Competing interests

The authors declare that they have no competing interests.

## Authors’ contributions

MBG and YYX conceived the study. MBG designed the project, processed the data, performed the analyses, interpreted the results, and wrote the manuscript. YYX was involved in project discussion. YYX and MBG provided financial support and reviewed the manuscript. All authors approved the manuscript.

## Acknowledgements

The results shown here are in whole or part based upon data generated by the TCGA Research Network: https://www.cancer.gov/tcga. The research has been supported by National Natural Science Foundation of China (NSFC) (Grant 31571350, U1611265, and 31871323) and Guangdong Basic and Applied Basic Research Foundation (2021A1515110972).

## Notes

### Competing Interest Statement

The authors have declared no competing interest.

