## Supplemental Info for "Pan-cancer RNA editing activity reveals complex editing functions and cancer immunotherapy biomarkers"

### Supplementary Information

#### Supplementary Results

##### Related publications for RNA editing associated RBPs with known relationship with DNA damage and repair

Published references of DNA damage and repair for RNA editing associated RBPs are as following: YTHDF2 [1], DDX51 [2], EWSR1 [3], TIA1 [4], STAU2 [5], and ZC3H11A [6]. In addition, DDX3X functions like RNaseH2 to perform ribonucleotide excision repair in R-loops to avoid genome instability and can also bind p53 to promote apoptosis upon DNA damage [7, 8].

#### Supplementary Figure and Legends

Figure. S1

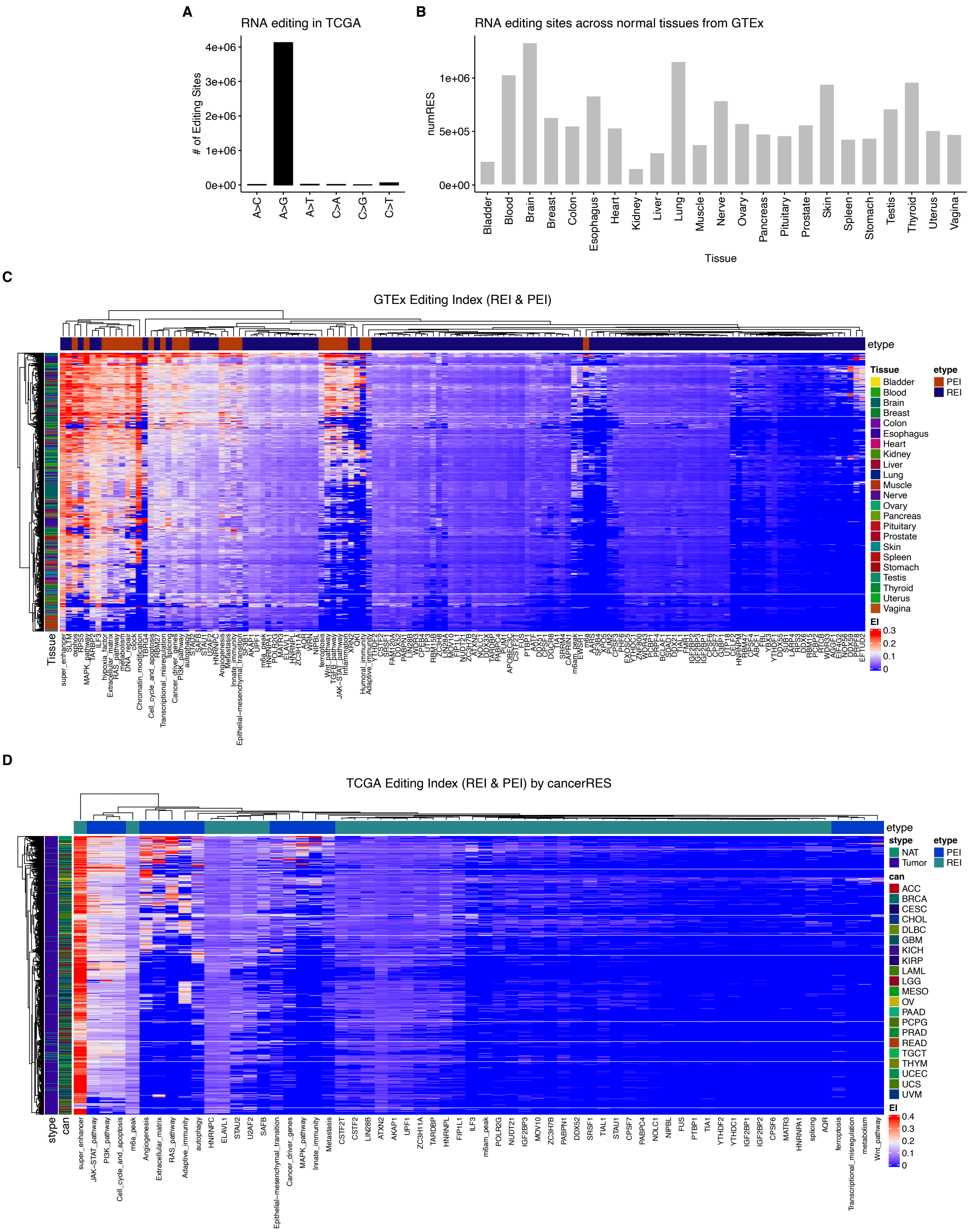


**Fig. S1, Charateristics of RES and EIs in TCGA and GTEx**. (**A**) Frequencies of six types of RESs in TCGA samples. (**B**) Number of RESs across tissues of GTEx. (**C**) All REIs and PEIs (colored heatmap values) calculated in this study for each sample from 23 tissues of GTEx. (**D**) All REIs and PEIs (colored heatmap values) calculated in this study for each sample from 21 TCGA cancer types, using cancerRES only. Rows are samples and columns are editing indexes (EIs). Sample types were also indicated.

Figure. S2

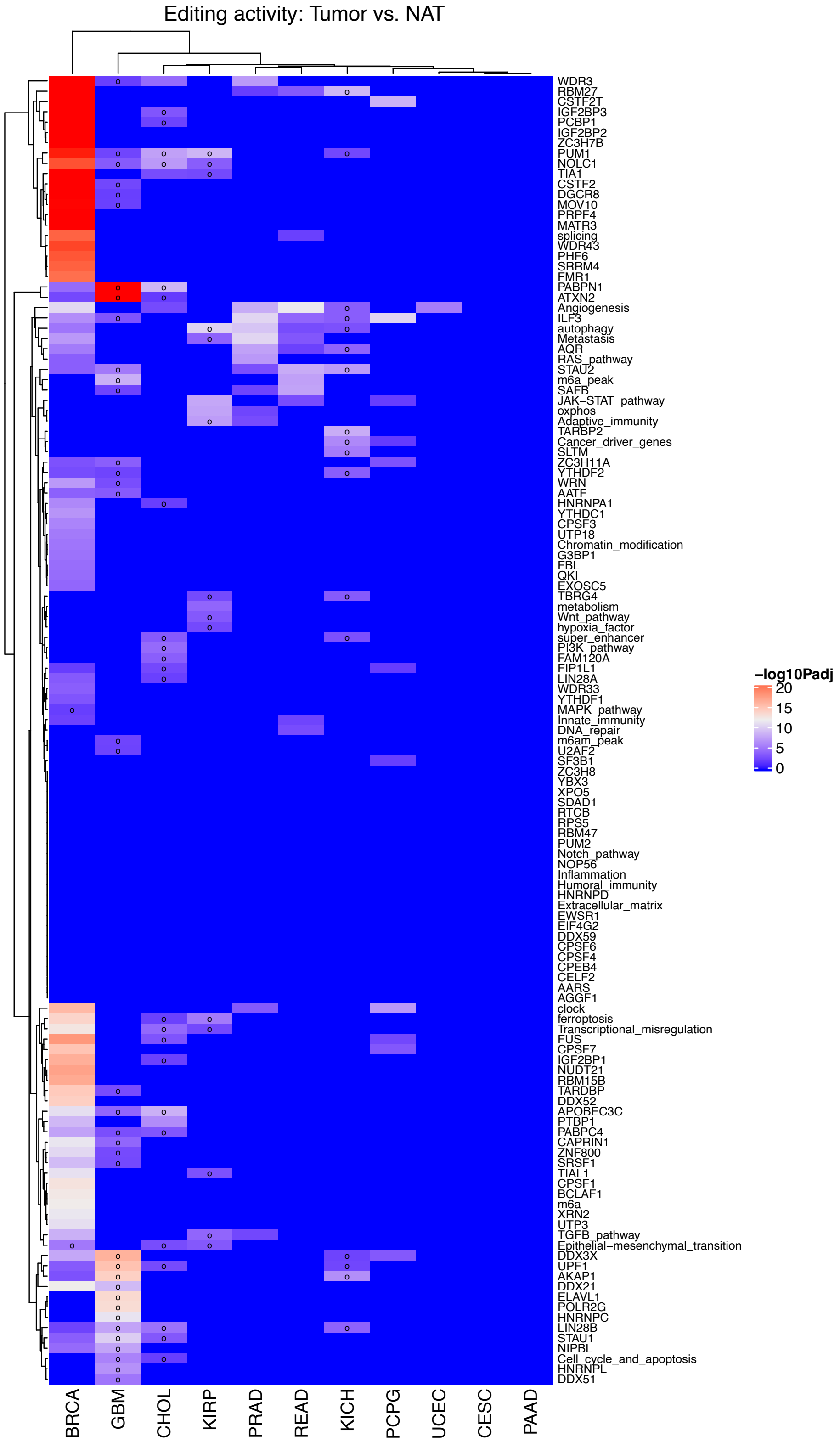


**Fig. S2, Differential EIs between tumor and adjacent normal samples**. Heatmap of differential REIs of RBPs comparing tumors with normal adjacent tissues (NATs). Colors stand for -log10 FDR values. Circles mark lower REIs in tumors than in NATs.

Figure. S3

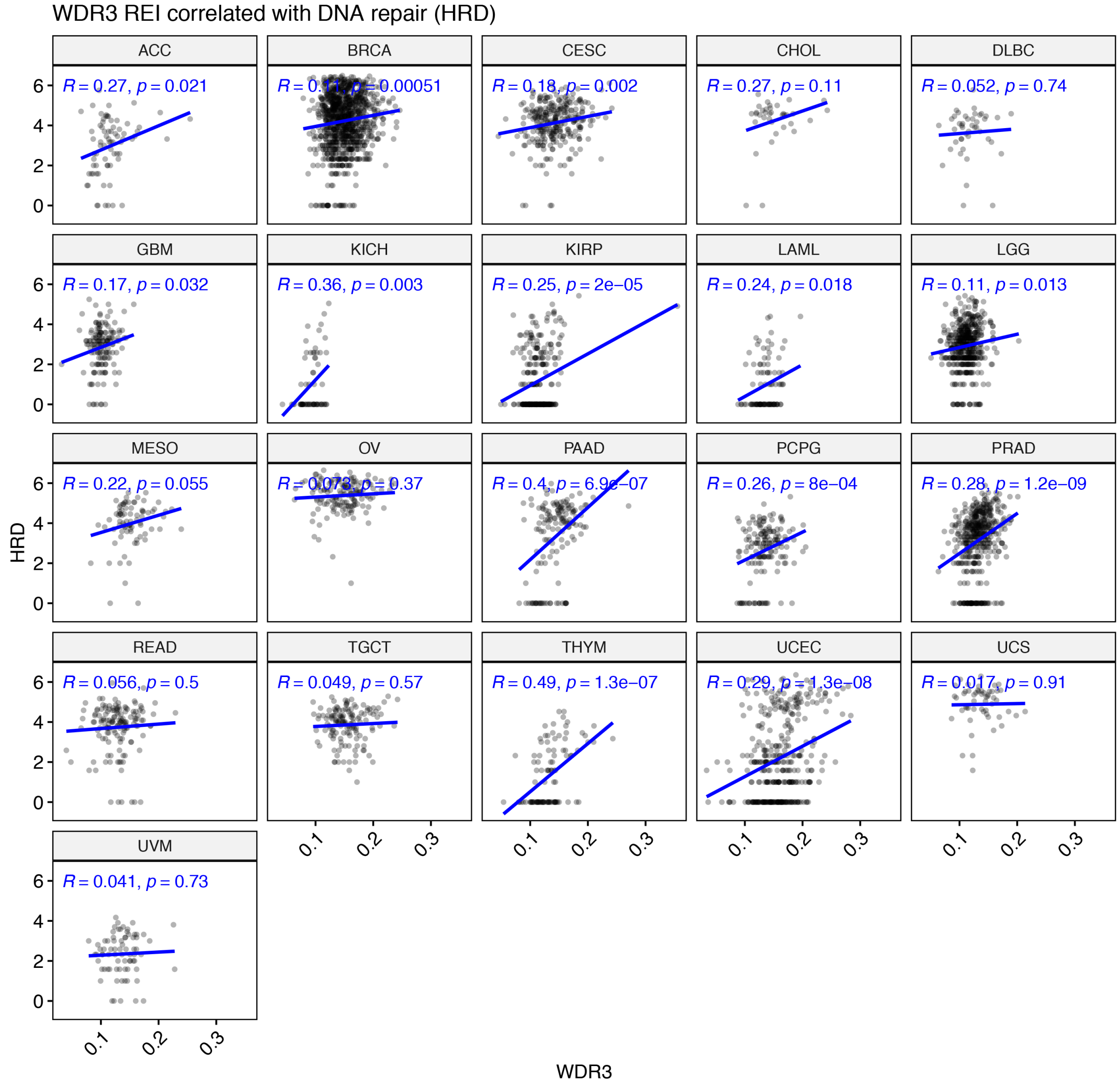


**Fig. S3, REI of WDR3 was correlated with DNA repair deficiency (HRD) in multiple cancer types**. Significant cancers included BRCA, KIRP, PAAD, PCPG, PRAD, THYM, and UCEC.

Figure. S4

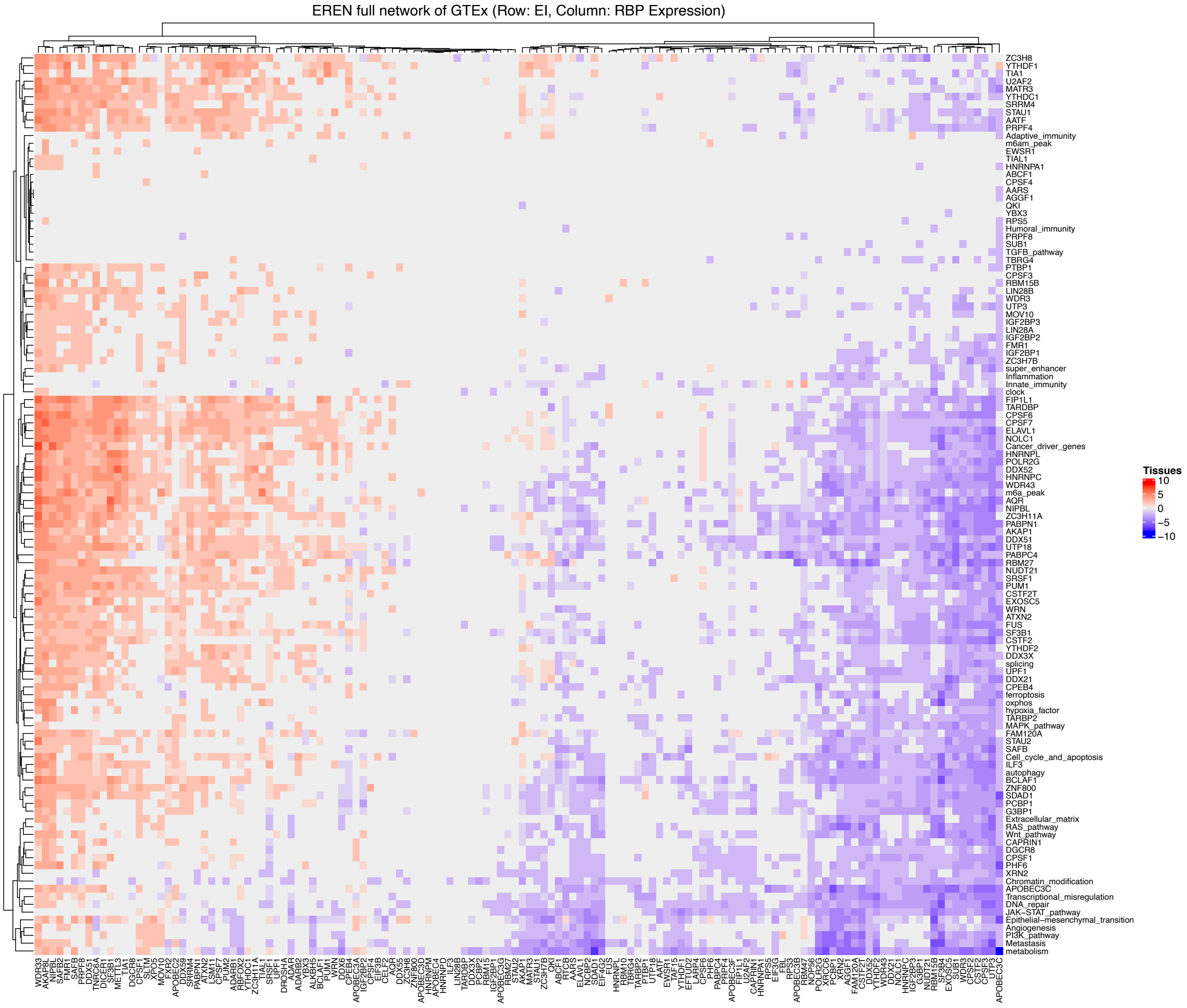


**Fig. S4, The full EREN network of GTEx**. Rows are EIs features and columns are RBP expression features. Colors represent tissue frequencies of each significant EI-RBP pair across GTEx tissues. Red stands for positive association and blue stands for negative association.

Figure. S5

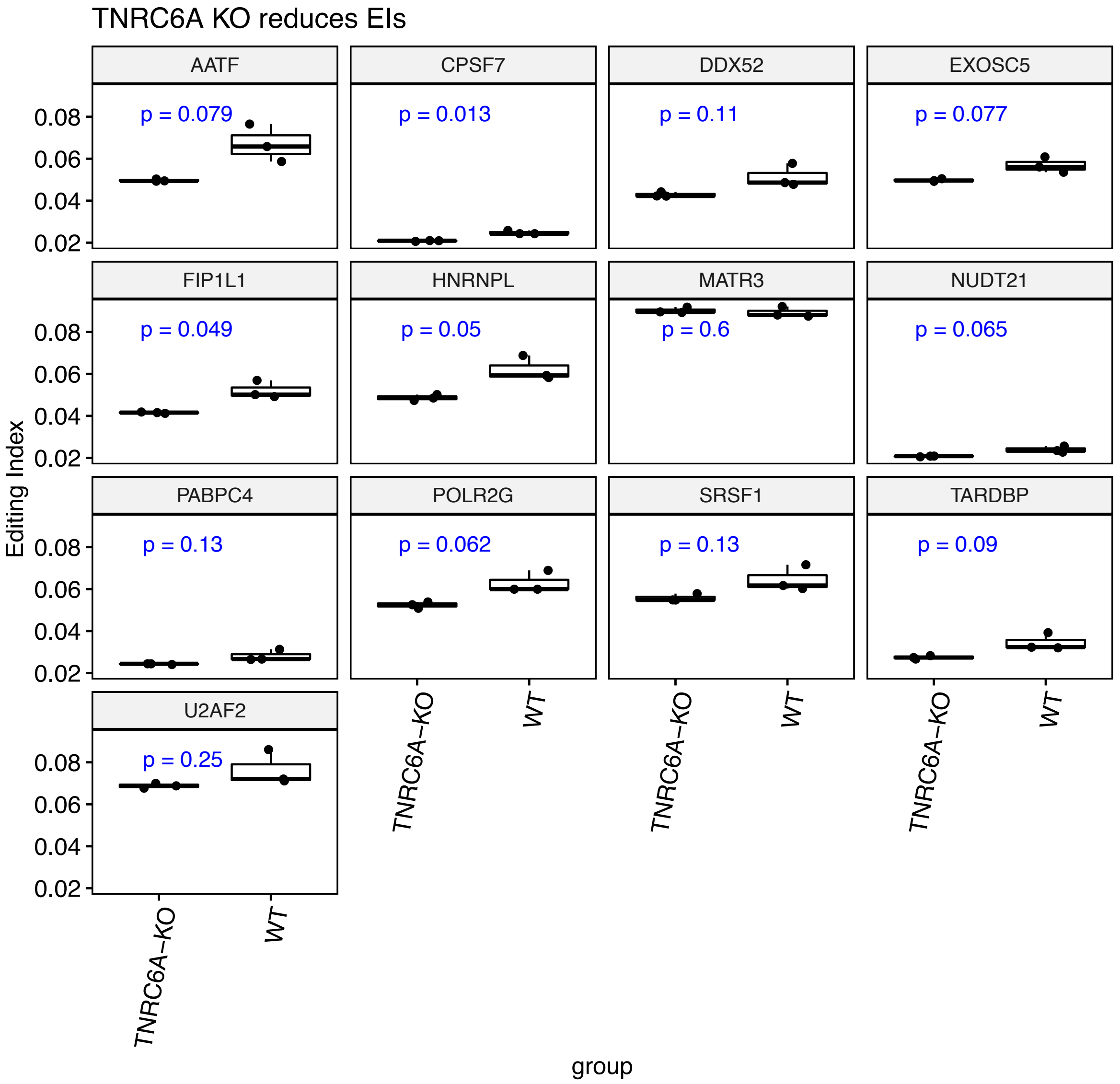


**Fig. S5, TNRC6A knockout (KO) reduced EIs of TNRC6A-associated RBPs (except MATR3) in the TCGA EREN core subnetwork**. Examples included EXOSC5, FIP1L1, HNRNPL, NUDT21, and POLR2G.

Figure. S6

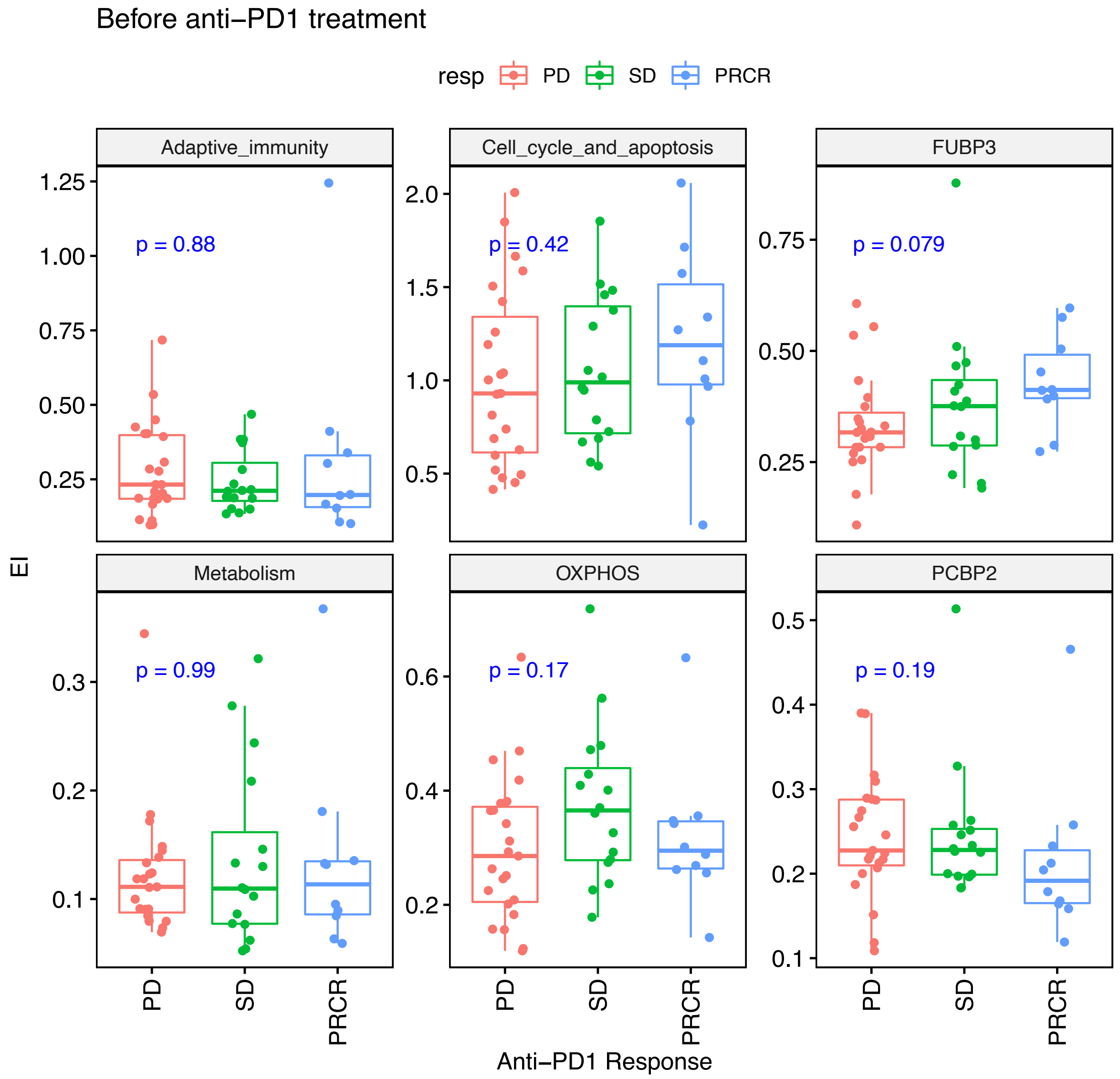


**Fig. S6, EI biomarkers during immunotherapy treatment cannot predict responses of anti-PD1 immunotherapy before treatment**. PD, progressive disease; SD, stable disease; PRCR, partial response or complete response.

**Supplementary Tables**

**Table S1**. Significant EI-Stemness pairs in TCGA tumor samples.

**Table S2**. Significant EI-TME pairs in TCGA tumor samples.
